# Dot1L licenses DNA demethylation to establish regulatory T cell identity

**DOI:** 10.64898/2025.12.03.692152

**Authors:** James J. Cameron, Tyler R. Colson, Xudong Li

## Abstract

Foxp3^+^ regulatory T (T_reg_) cells rely on DNA demethylation to establish and maintain the gene expression program that defines their identity and suppressive function. Although sustained transcription of Treg-specific genes is known to drive this demethylation, the precise mechanism has remained unclear. Here, we show that Dot1L-catalyzed methylation of histone H3 at lysine 79 (H3K79me) is essential for Treg-specific DNA demethylation in both thymic and induced Treg lineages. H3K79me promotes chromatin activation and recruits TET family DNA demethylases to key regulatory loci. Treg-restricted deletion of Dot1L disrupts locus-specific DNA demethylation, diminishes expression of core Treg genes, and precipitates a fatal, early-onset autoimmune syndrome. Conversely, pharmacologic enhancement of TET activity during differentiation rescues DNA demethylation, restores Treg-gene expression, and reinstates suppressive function even in the presence of Dot1L inhibition. Together, these findings identify a critical epigenetic axis—Dot1L-mediated H3K79 methylation driving TET-dependent DNA demethylation—that safeguards Treg cellular identity, function, and immune homeostasis.

## Introduction

Regulatory T cells (Tregs) are a specialized subset of CD4⁺ T cells that enforce self-tolerance and preserve immune homeostasis (1, 2). Their development and function depend on sustained expression of the X-linked transcription factor Foxp3, which both initiates Treg lineage commitment and maintains suppressive activity in mature cells (3–7). Fate-mapping studies have demonstrated that Foxp3 expression in committed Tregs remains remarkably stable at steady state and during inflammation in vivo (8, 9). However, a small fraction of Tregs can lose Foxp3 expression in inflamed tissues—becoming “exTregs” that secrete pro-inflammatory cytokines and exacerbate autoimmunity (10–12). Despite the critical importance of Foxp3 stability for Treg function, the molecular mechanisms that establish and preserve the epigenetic memory of Foxp3 expression remain incompletely defined, posing a major barrier to the development of robust Treg-based cellular therapies (13, 14).

Epigenetic regulation—through DNA methylation and histone modifications—is essential for establishing the Treg-specific gene program (4, 15, 16). During Treg differentiation, key regulatory loci become demethylated at Treg-specific demethylated regions (TSDRs), most notably the conserved non-coding sequence 2 (CNS2) within the Foxp3 locus (4, 15, 17). In mature Tregs, CNS2 remains stably demethylated, a feature required for long-term Foxp3 expression, whereas deletion of CNS2 precipitates Treg instability and autoimmune pathology (18, 19). By contrast, induced Tregs (iTregs) generated in vitro from naïve CD4⁺ T cells under TGF-β and IL-2 conditions largely retain CNS2 methylation, which correlates with their propensity to lose Foxp3 and suppressive function under inflammatory stress (15). Beyond Foxp3 CNS2, demethylation at multiple additional TSDRs is associated with the activation of genes critical for Treg identity and function (4). Together, these observations underscore the importance of finely tuned DNA demethylation in sustaining Treg lineage stability and immunosuppressive capacity.

Our previous work revealed that sustained transcriptional activation of *Foxp3*—rather than Foxp3 protein expression—drives CNS2 demethylation in iTregs (20). This mechanism ensures that stable Foxp3 expression and Treg lineage commitment via CNS2 demethylation occur only in cells receiving continuous instructive signals for *Foxp3* transcription. Building on this insight, we recently developed an approach to epigenetically reprogram autoreactive pro-inflammatory effector T (Teff) cells into Tregs with a superior ability to control established autoimmune inflammation compared with iTregs and endogenous Tregs (21). However, the molecular link between persistent Foxp3 transcription and CNS2 demethylation remains unknown, limiting further optimization of this therapeutic strategy aimed at improving reprogramming efficiency. In particular, although TET family DNA demethylases and DNA methyltransferases (DNMTs) are key regulators of DNA methylation at the *Foxp3* locus (22–25), it is unclear how *Foxp3* transcription influences the recruitment or activity of these enzymes to achieve demethylation. Moreover, it remains to be determined whether this transcription-induced demethylation mechanism similarly governs other Treg-specific demethylated regions.

Gene transcription is regulated by multiple histone modifications, including methylation of histone H3 at lysine 79 (H3K79), which is enriched within the bodies of actively transcribed genes (26–28)—coincident with many Treg-specific demethylated regions (TSDRs) (4, 29). Because H3K79 methylation correlates strongly with gene activity, we investigated the role of Dot1L, the sole H3K79 methyltransferase, in directing TSDR demethylation and establishing Treg identity. We found that Dot1L-dependent H3K79 methylation is essential for demethylation of CNS2 and other TSDRs in both induced and thymic Treg lineages. Inhibition of Dot1L reduced chromatin accessibility and TET demethylase binding at CNS2 and other Foxp3 regulatory elements, whereas CRISPR/Cas9-driven chromatin activation of CNS2 or TET enhancement with vitamin C restored both DNA demethylation and Foxp3 stability despite Dot1L blockade. Consistent with these in vitro findings, Treg-specific deletion of Dot1L in mice led to fatal inflammatory disease owing to defective Treg gene-program establishment. Moreover, enhancing TET activity during in vitro differentiation of tTreg precursors rescued Treg function when Dot1L was inhibited. Together, these results demonstrate that Dot1L-mediated H3K79 methylation drives Treg-specific DNA demethylation and is indispensable for maintaining Treg identity and suppressive function.

## Results

### Dot1L Is Required for CNS2 Demethylation and Foxp3 Stability in iTregs

To determine whether H3K79 methylation accompanies *Foxp3* induction during Treg differentiation, we performed chromatin immunoprecipitation (ChIP) for H3K79me2 at the *Foxp3* locus in Foxp3⁺ induced Tregs (iTregs) and Foxp3⁻ Th0 controls. iTregs showed marked enrichment of H3K79me2 at both CNS1, the enhancer critical for *Foxp3* induction (17), and CNS2, compared with Th0 cells (**Fig. 1A**). Treatment of differentiating iTregs with the Dot1L inhibitor SGC0946 (SGC) (30) substantially reduced H3K79me2 levels at CNS1, CNS2, and the *Foxp3* transcription start site (TSS) (**Fig. 1B**), confirming that Dot1L mediates H3K79 methylation at these key regulatory elements.

**Figure 1.**
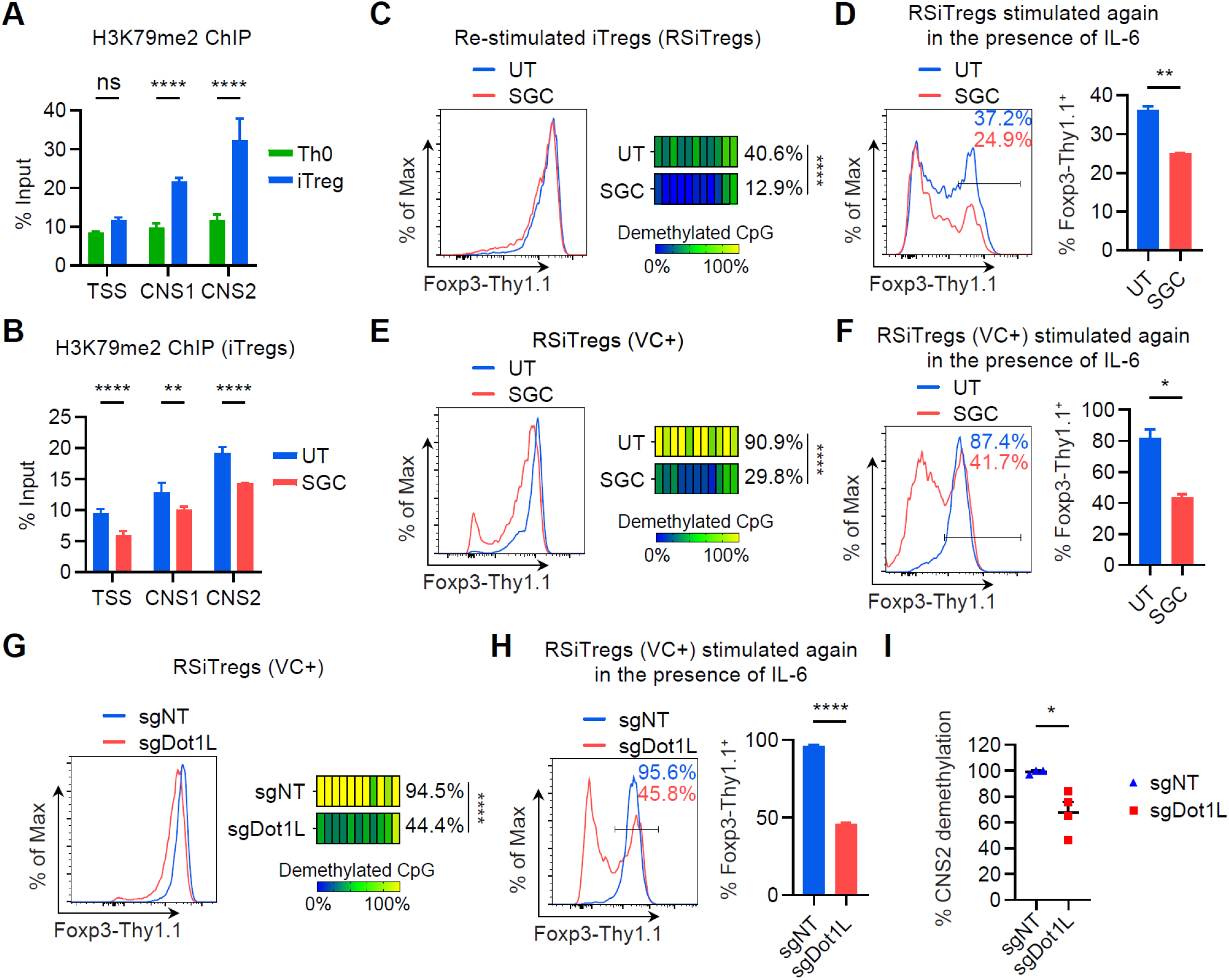
Dot1L is indispensable for CNS2 demethylation and stable *Foxp3* expression in iTregs. (A) Normalized H3K79me2 levels measured by ChIP-qPCR at *Foxp3* TSS, CNS1, and CNS2 in Th0 and iTregs generated by stimulating CD4^+^Foxp3-Thy1.1^−^CD44^lo^CD62L^hi^ Tn cells from *Foxp3^Thy1.1^* mice with plate-coated anti-CD3/CD28 antibodies in the presence of IL-2 alone (Th0) or IL-2 and TGFβ (iTregs) for 4 days. (B) Normalized H3K79me2 levels measured by ChIP-qPCR at *Foxp3* TSS, CNS1, and CNS2 in *Foxp3^Thy1.1^* re-stimulated iTregs (RSiTregs) differentiated in the presence or absence of 2 µM of the Dot1L inhibitor SGC0946 (SGC). (C) Flow cytometry of Foxp3-Thy1.1 expression in *Foxp3^Thy1.1^* RSiTregs differentiated in the presence or absence of 2 µM of SGC. Heatmaps show DNA demethylation status of *Foxp3* CNS2 analyzed with bisulfite-sequencing. Each bar represents a CpG site. (D) Flow cytometry of Foxp3-Thy1.1 expression in *Foxp3^Thy1.1^* RSiTregs differentiated in the presence or absence of 2 µM of SGC and re-stimulated again with anti-CD3/CD28 beads for 3 days in the presence of 100 U/ml of IL-2 and 20 ng/ml of IL-6. (E) Flow cytometry of Foxp3-Thy1.1 expression in *Foxp3^Thy1.1^* RSiTregs differentiated in the presence of 1 µg/ml of vitamin C (VC) and with or without 2 µM of SGC. Heatmaps show DNA demethylation status of *Foxp3* CNS2. (F) Flow cytometry of Foxp3-Thy1.1 expression in *Foxp3^Thy1.1^* RSiTregs differentiated in the presence of 1 µg/ml of VC, with or without 2 µM of SGC, and re-stimulated again in the presence of 100 U/ml of IL-2 and 20 ng/ml of IL-6. (G) Flow cytometry of Foxp3-Thy1.1 expression in *Foxp3^Thy1.1^R26^Cas9^* RSiTregs transduced with a retroviral vector (RV) expressing a single guide RNA targeting Dot1L (sgDot1L) or a non-targeting sgRNA (sgNT). Heatmaps show DNA demethylation status of *Foxp3* CNS2. (H) Flow cytometry of Foxp3-Thy1.1 expression in *Foxp3^Thy1.1^R26^Cas9^* RSiTregs transduced with the sgDot1L-RV or the sgNT-RV and re-stimulated again in the presence of 100 U/ml of IL-2 and 20 ng/ml of IL-6. (I) Percentages of CNS2 demethylation in *Foxp3^Thy1.1^* iTregs that were doubly transduced with the Cas9-RV and the sgDot1L-RV or the sgNT-RV, adoptively co-transfer with CD45 congenically distinct *Foxp3^Thy1.1^* Tn cells into *Rag1^−/−^* mice, and sort-purified 5 weeks later. Each point represents % CNS2 demethylation in individual mouse. Mean ± SEM. *p < 0.05, **p < 0.01, ****p < 0.0001, one-way ANOVA and Holm-Šídák test in (A and B), unpaired two-sided t-test in (D, F, H, and I), and two-way ANOVA in (C, E, and G).

Next, we asked whether Dot1L-dependent H3K79 methylation is necessary for CNS2 demethylation. Re-stimulation of Foxp3^Thy1.1^ iTregs to sustain *Foxp3* transcription enhances CNS2 demethylation; however, inclusion of SGC during re-stimulation reduced CNS2 demethylation by over threefold without altering Thy1.1 reporter levels (**Fig. 1C**). When these re-stimulated iTregs (RSiTregs) were challenged with IL-6—which destabilizes Foxp3 in the absence of a demethylated CNS2 (18, 19)—we observed significantly decreased *Foxp3* stability in SGC-treated cells (**Fig. 1D**). Thus, while Dot1L inhibition has minimal effect on initial *Foxp3* induction, it critically impairs CNS2 demethylation and compromises Foxp3 maintenance.

Given the critical role of TET proteins in mediating CNS2 demethylation (22–24), we asked whether Dot1L supports TET-dependent demethylation in iTregs. Addition of the TET activator vitamin C (VC) (31), during iTreg restimulation increased CNS2 demethylation in RSiTregs from approximately 40% to over 90% (**Fig. 1E**). By contrast, pharmacologic inhibition of Dot1L during restimulation reduced CNS2 demethylation from 90% to 30% and markedly impaired the stability of Foxp3 expression (**Fig. 1, E and 1F**). These data indicate that Dot1L-mediated H3K79 methylation promotes TET-dependent CNS2 demethylation and thereby stabilizes Foxp3 expression in iTregs.

To validate these findings genetically, we transduced *Foxp3^Thy1.1^R26^Cas9^*iTregs with either a Dot1L-targeting sgRNA (sgDot1L) or a non-targeting control (sgNT). Upon re-stimulation in VC, Dot1L-deficient RSiTregs displayed only a slight reduction in Thy1.1 expression but showed a pronounced loss of CNS2 demethylation and stability of *Foxp3* expression relative to sgNT controls (**Fig. 1G** and **1H**). In vivo, co-transfer of sgDot1L-or sgNT-transduced iTregs into *Rag1^−/−^* mice alongside naïve CD4⁺ T cells revealed significantly reduced CNS2 demethylation in Dot1L-deficient donor cells five weeks post-transfer (**Fig. 1I**). Collectively, these experiments demonstrate that Dot1L-mediated H3K79 methylation is essential for CNS2 demethylation and *Foxp3* stability in iTregs both in vitro and in vivo.

### Dot1L Facilitates Chromatin Activation to Enable CNS2 Demethylation

We next investigated how H3K79 methylation influences CNS2 chromatin state. Inhibition of Dot1L in iTregs led to pronounced reductions in the active histone marks H3K27ac and H3K9ac at CNS2 and other Foxp3 cis-elements (**Fig. 2, A and B**), accompanied by an increase in the repressive mark H3K27me3 (**Fig. 2C**). H3K4me3 at the promoter remained unchanged, whereas H3K4me1—associated with poised enhancers—was elevated upon Dot1L inhibition (**Fig. 2, D and E**). These data indicate that Dot1L-mediated H3K79 methylation promotes an open, active chromatin environment at CNS2.

**Figure 2.**
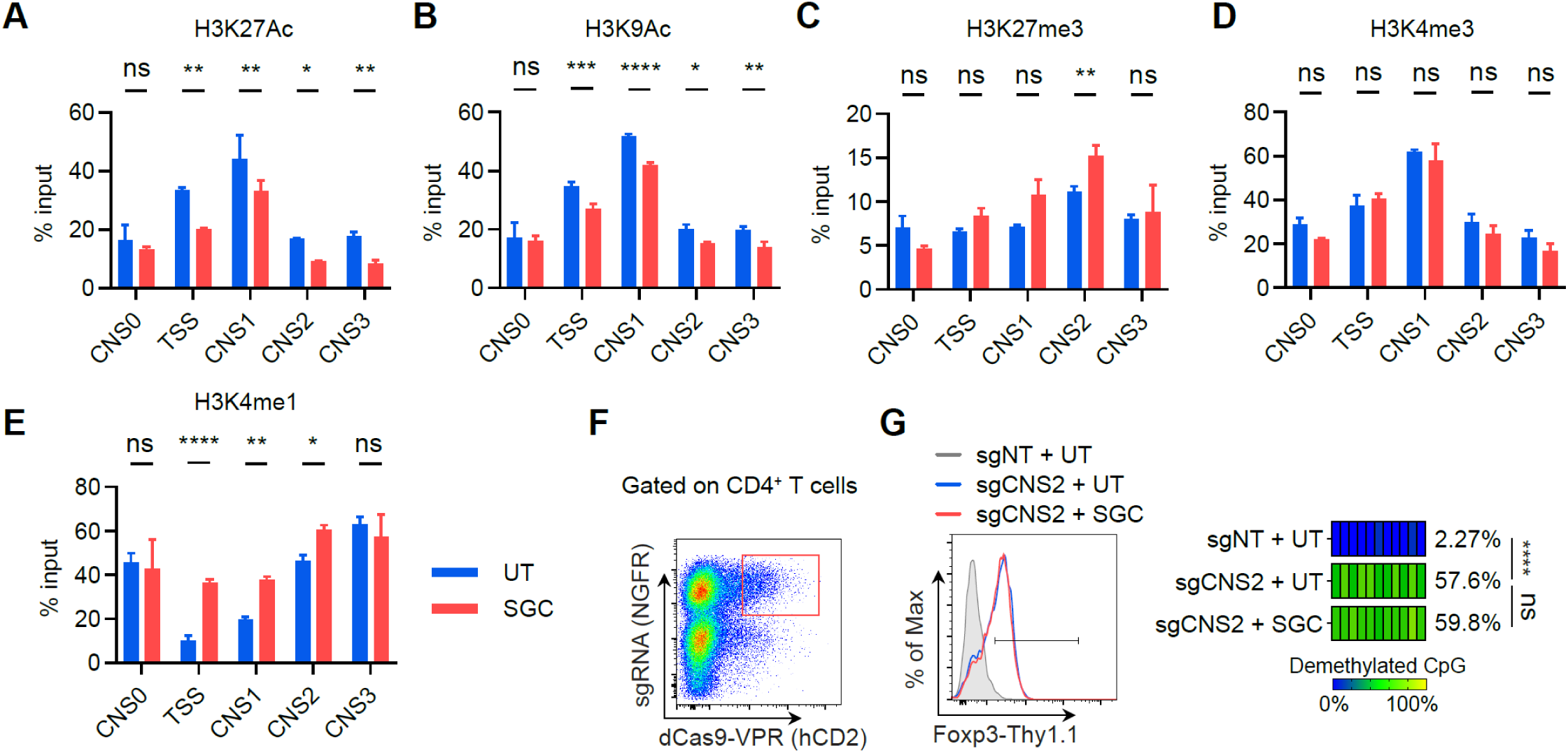
Dot1L facilitates *Foxp3* chromatin activation to promote CNS2 demethylation. (A-E) Normalized levels of H3K27Ac (A), H3K9Ac (B), H3K27me3 (C), H3K4me3 (D), and H3K4me1 (E) measured by ChIP-qPCR at *Foxp3* CNS0, TSS, CNS1, CNS2, and CNS3 in RSiTregs differentiated in the presence or absence of 2 µM of SGC. (F-G) Flow cytometry (F) of the expression of the sgRNA-RV marker NGFR and the dCas9-VPR-RV marker hCD2 in *Foxp3^Thy1.1^* Tn cells stimulated with plate-coated anti-CD3/CD28 antibodies in the presence of 1000 U/ml of IL-2 and 1 µg/ml of VC, and transduced with the RV expressing sgCNS2 and the RV expressing dCas9-VPR. Flow cytometry (G) of Foxp3-Thy1.1 expression in NGFR^+^hCD2^+^CD4^+^ T cells as gated in (F) after TCR-stimulation in the presence of IL-2, VC, and with or without SGC. Heatmaps in (G) show DNA demethylation status of *Foxp3* CNS2. Mean ± SEM. *p < 0.05, **p < 0.01, ***p < 0.001, ****p < 0.0001, one-way ANOVA and Holm-Šídák test in (A-E) and two-way ANOVA in (G).

To test whether chromatin activation alone can drive CNS2 demethylation independent of Dot1L, we targeted a synthetic CRISPR/dCas9-based transcriptional activator (dCas9-VPR) to CNS2 in naïve T cells (32). In the presence of sgCNS2, dCas9-VPR induced robust *Foxp3* expression and CNS2 demethylation, even when Dot1L was inhibited by SGC (**Fig. 2, F and G**). Thus, enhanced chromatin activation can bypass the need for H3K79 methylation to confer CNS2 demethylation.

### Dot1L Promotes TET Recruitment to Enhance CNS2 Demethylation

Given the established role of TET proteins in CNS2 demethylation and our findings implicating Dot1L in TET-dependent demethylation (**Fig. 1E**), we next asked whether Dot1L-mediated H3K79 methylation influences TET protein binding at CNS2. Strikingly, inhibition of Dot1L with SGC during iTreg differentiation significantly diminished Tet2 and Tet3 binding across most known conserved *Foxp3* cis-regulatory elements (**Fig. 3, A and B**), suggesting that Dot1L is critical for recruiting TET proteins to CNS2. To determine whether this impaired TET recruitment underlies the reduced CNS2 demethylation in SGC-treated iTregs, we tested whether increasing the TET activator VC from 2 µg/ml to 10 µg/ml could rescue demethylation. Notably, higher VC concentrations fully restored CNS2 demethylation even in the presence of SGC (**Fig. 3C**), accompanied by a recovery of Foxp3 expression stability (**Fig. 3D**). Together, these results demonstrate that Dot1L-dependent H3K79 methylation facilitates CNS2 demethylation, at least in part by enhancing the recruitment of TET demethylases.

**Figure 3.**
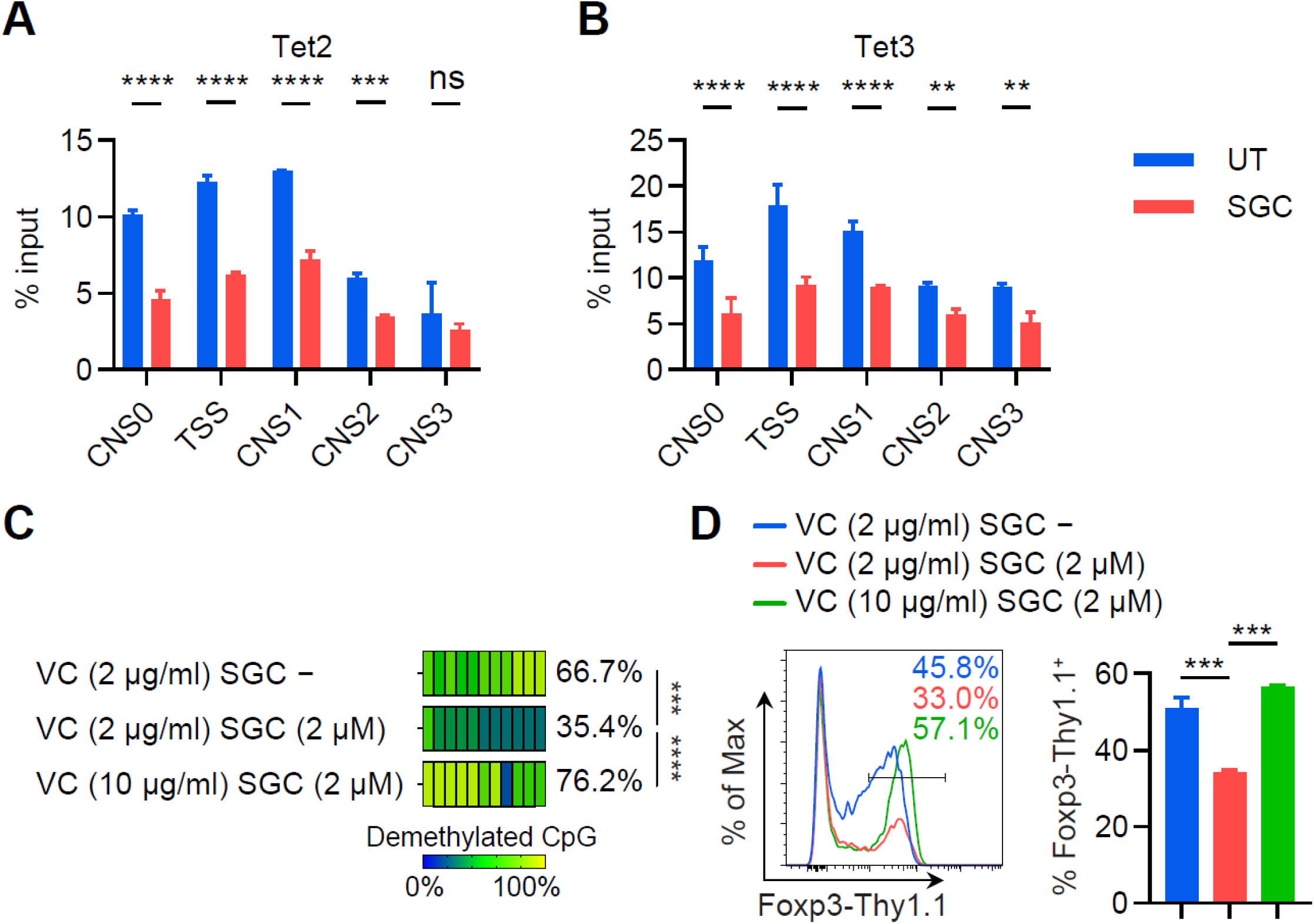
Dot1L promotes CNS2 demethylation by enhancing TET recrutiment. (A,B) Normalized Tet2 (A) and Tet3 (B) binding measured by ChIP-qPCR at *Foxp3* CNS0, TSS, CNS1, CNS2, and CNS3 in RSiTregs differentiated in the presence or absence of 2 µM of SGC. (C,D) Heatmaps (C) of DNA demethylation status of *Foxp3* CNS2 in *Foxp3^Thy1.1^* iTregs differentiated in the presence of 2 or 10 µg/ml of VC and with or without 2 µM of SGC. Flow cytometry (D) of Foxp3-Thy1.1 expression in iTregs re-stimulated in the presence of 100 U/ml of IL-2 and 20 ng/ml of IL-6. Mean ± SEM. **p < 0.01, ***p < 0.001, ****p < 0.0001, one-way ANOVA and Holm-Šídák test in (A, B, and D) and two-way ANOVA in (C).

### Treg-Specific Dot1L Deletion Impairs tTreg TSDR Demethylation In Vivo

To assess the role of Dot1L in TSDR demethylation in tTregs in vivo, we generated female *Foxp3^Cre-YFP/+^Dot1L^fl/fl^* mice and leveraged their random inactivation of the X-linked *Foxp3* allele. In these mice, Dot1L-deficient (YFP^+^) and Dot1L-sufficient (YFP^−^) Tregs coexist, avoiding confounding effects from systemic autoimmunity that could arise from Treg-specific Dot1L deletion. Control *Foxp3^Cre-YFP/+^Dot1L^fll+^* littermates provided Dot1L-sufficient YFP^+^ Tregs for comparison. Both cohorts displayed similar frequencies of YFP^+^CD25^+^ Tregs in the thymus (**Fig. 4A**). However, Dot1L-deficient mice exhibited a trending reduction of the frequencies of mature (CD24^lo^CD73^hi^) tTregs (**Fig. 4B**), and those mature tTregs displayed significantly reduced DNA demethylation across most TSDRs compared to controls (**Fig. 4C**). These data demonstrate that Dot1L is essential for TSDR demethylation during thymic Treg development.

**Figure 4.**
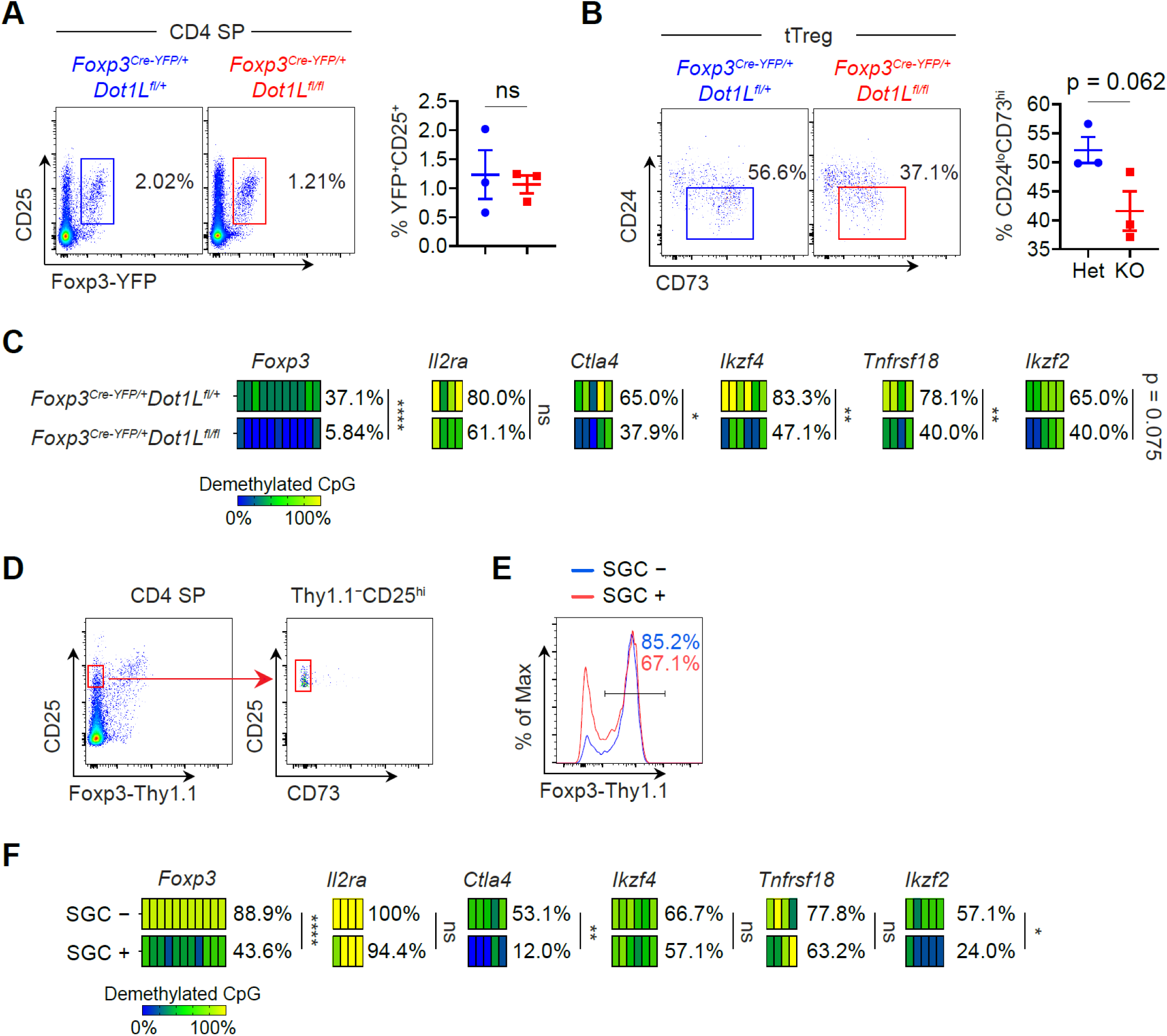
Treg-specific Dot1L deletion impairs DNA demethylation in thymic Tregs. (A) Flow cytometry of Foxp3-YFP and CD25 expression in CD4SP thymocytes from female *Foxp3*^Cre-YFP/+^*_Dot1L_^fl/+^* and Foxp3^Cre-YFP/+^Dot1L*^fl/fl^* mice. (B) Flow cytometry of CD73 and CD24 expression in Foxp3-YFP^+^CD25^+^ thymic Tregs from female *Foxp3^Cre-YFP/+^Dot1L^fl/+^* and *Foxp3^Cre-YFP/+^Dot1L^fl/fl^* mice. (C) Heatmaps of DNA demethylation status of Tregs-specific demethylation regions (TSDRs) in Foxp3-YFP^+^CD25^+^CD24^lo^CD73^hi^ mature thymic Tregs from female *Foxp3^Cre-YFP/+^Dot1L^fl/+^* and *Foxp3^Cre-YFP/+^Dot1L^fl/fl^* mice. (D-F) Flow cytometry (D) of CD25 and Foxp3-Thy1.1 expression (left) in CD4SP thymocytes and CD25 and CD73 expression (right) in Foxp3-YFP^−^CD25^+^ CD4SP thymocytes from 11-day-old *Foxp3^Thy1.1^* mice to show the gating strategy for tTreg precursors. Flow cytometry (E) of Foxp3-Thy1.1 expression in *Foxp3^Thy1.1^* tTreg precursors stimulated with anti-CD3/CD28 beads in the presence of IL-2 and with or without 2 µM of SGC. Heatmaps in (F) show DNA demethylation status of *Foxp3* CNS2 and *Ctla4* TSDR in tTreg precursors (Pre-Treg) prior to in vitro TCR stimulation or after TCR stimulation in the presence or absence of SGC. Mean ± SEM. *p < 0.05, **p < 0.01, ***p < 0.001, ****p < 0.0001, two-way ANOVA in (C and E).

To determine whether Dot1L’s histone methyltransferase activity is required for TSDR demethylation in tTregs, we isolated Foxp3-Thy1.1^−^CD25^hi^CD73^lo^ tTreg precursors (Pre-tTregs) from *Foxp3^Thy1.1^* mice—using CD73 to distinguish developing thymocytes from recirculating mature T cells (33) (**Fig. 4D**)—and stimulated them in vitro with anti-CD3/CD28 plus IL-2 in the presence or absence of SGC. Although SGC only modestly decreased Foxp3-Thy1.1^+^ cell frequency (**Fig. 4E**), it significantly impaired demethylation at *Foxp3* CNS2 and *Ctla4* and *Ikzf2* TSDRs. In contrast, demethylation at *Ikzf4* and *Tnfrsf18* TSDRs was unaffected (**Fig. 4F**), unlike the reduction observed with genetic Dot1L deletion (**Fig. 4C**). This discrepancy suggests that either residual H3K79 methylation in precursors suffices for demethylation at these sites or that Dot1L can facilitate demethylation via H3K79-independent mechanisms. Collectively, these findings indicate that Dot1L-mediated H3K79 methylation is essential for optimal TSDR demethylation in most loci during tTreg development.

### Treg-specific Deletion of Dot1L in Mice Resulted in Fatal Systemic Autoimmunity

To investigate the physiological consequences of Dot1L ablation in Tregs, we generated male *Foxp3^Cre-YFP/y^Dot1L^fl/fl^*mice (with Dot1L-deficient Tregs) and control *Foxp3^Cre-YFP/y^Dot1L^fl/+ or^ ^+/+^*littermates. *Foxp3^Cre-YFP/y^Dot1L^fl/fl^* mice showed severe growth retardation (**Fig. 5A**) and typically succumbed to disease between 3-5 weeks of age. These mice developed a profound lymphoproliferative syndrome, characterized by splenomegaly and markedly increased CD8^+^ T cell counts in the spleen, as well as elevated total cellularity and expanded CD4^+^ and CD8^+^ T cell populations in peripheral lymph nodes (**Fig. 5, B and E**). Moreover, both CD4^+^ and CD8^+^ T cells in these mice displayed an activated (CD62L^lo^CD44^hi^) phenotype in spleen and lymph nodes (**Fig. 5, F and G**). Furthermore, CD4^+^ T cells from mutant mice showed markedly increased production of IFNγ, IL-4, and IL-17A (**Fig. 5H**), indicating global failure of immune regulation by Dot1L-deficient Tregs. Collectively, these findings suggest that Dot1L-dependent TSDR demethylation is essential for maintaining Treg suppressor function, and its loss precipitates lethal, early-onset autoimmunity.

**Figure 5.**
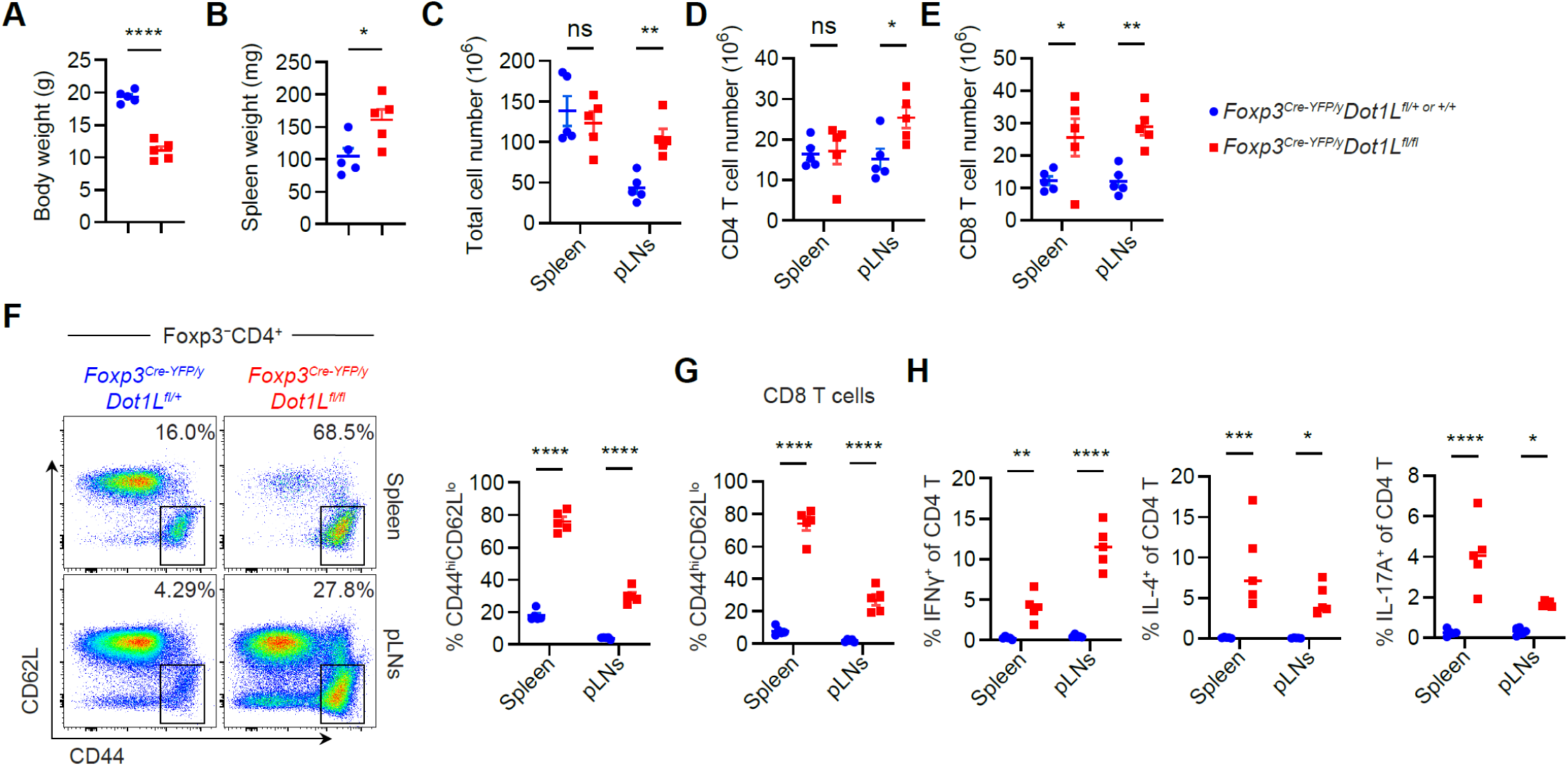
Mice with Treg-specific Dot1L deletion develop fatal inflammatory disorder. (A,B) Body weight (A) and spleen weight (B) of 3-to 5-week-old male *Foxp3^Cre-YFP/y^Dot1L^fl/fl^* and *Foxp3^Cre-YFP/y^Dot1L^fll+ or +/+^* littermate control mice. n = 5. (C-E) Cellularity (C), CD4^+^ T cell numbers (C), and CD8^+^ T cell numbers (D) in spleens and peripheral lymph nodes (pLNs) from 3-to 5-week-old male *Foxp3^Cre-YFP/y^Dot1L^fl/fl^* and *Foxp3^Cre-YFP/y^Dot1L^fll+ or +/+^* littermate control mice. n = 5. (F) Flow cytometry (left) of CD44 and CD62L and frequencies (right) of CD44^hi^CD62L^lo^ cells in Foxp3-Thy1.1^−^CD4^+^ T cells in spleens and pLNs from 3-to 5-week-old male *Foxp3^Cre-YFP/y^Dot1L^fl/fl^* and *Foxp3^Cre-YFP/y^Dot1L^fll+ or +/+^* littermate control mice. n = 5. (G) Frequencies of CD44^hi^CD62L^lo^ cells in CD8^+^ T cells in spleens and pLNs from 3-to 5-week-old male *Foxp3^Cre-YFP/y^Dot1L^fl/fl^* and *Foxp3^Cre-YFP/y^Dot1L^fll+ or +/+^* littermate control mice. n = 5. (H) Frequencies of IFNγ^+^, IL-4^+^, and IL-17A^+^ cells in CD4^+^ T cells in spleens and pLNs from 3-to 5-week-old male *Foxp3^Cre-YFP/y^Dot1L^fl/fl^* and *Foxp3^Cre-YFP/y^Dot1L^fll+ or +/+^* littermate control mice. n = 5. Mean ± SEM. *p < 0.05, **p < 0.01, ***p < 0.001, ****p < 0.0001, unpaired two-sided t-test in (A, B) and one-way ANOVA and Holm-Šídák test in (C-H).

### Treg-specific Dot1L Deletion Impairs Expression of Treg Identity Genes

To elucidate how Dot1L governs Treg identity and function, we first assessed Treg frequencies and phenotype in mice with Treg-specific Dot1L deletion. Similar to CNS2-deficient mice, Treg-specific Dot1L knockout mice showed reduced splenic (but not lymph node) Treg frequencies (**Fig. 6A**), suggesting defective Treg homeostasis contributes to the observed inflammatory phenotype. Moreover, Dot1L-deficient (YFP^+^) Tregs in female *Foxp3^Cre-YFP/+^Dot1L^fl/fl^* mice exhibited markedly lower surface expression of key suppressive receptors (ICOS, TIGIT, Nrp1) and reduced levels of multiple TSDR-linked proteins—Foxp3, CD25, CTLA-4, EOS, GITR, and Helios (4)— relative to Dot1L-sufficient (YFP^+^) Tregs from *Foxp3^Cre-YFP/+^Dot1L^fl/fl^*mice (**Fig. 6, B and C**). These data indicate that Dot1L is essential for robust expression of molecules critical to Treg lineage stability and suppressive function.

**Figure 6.**
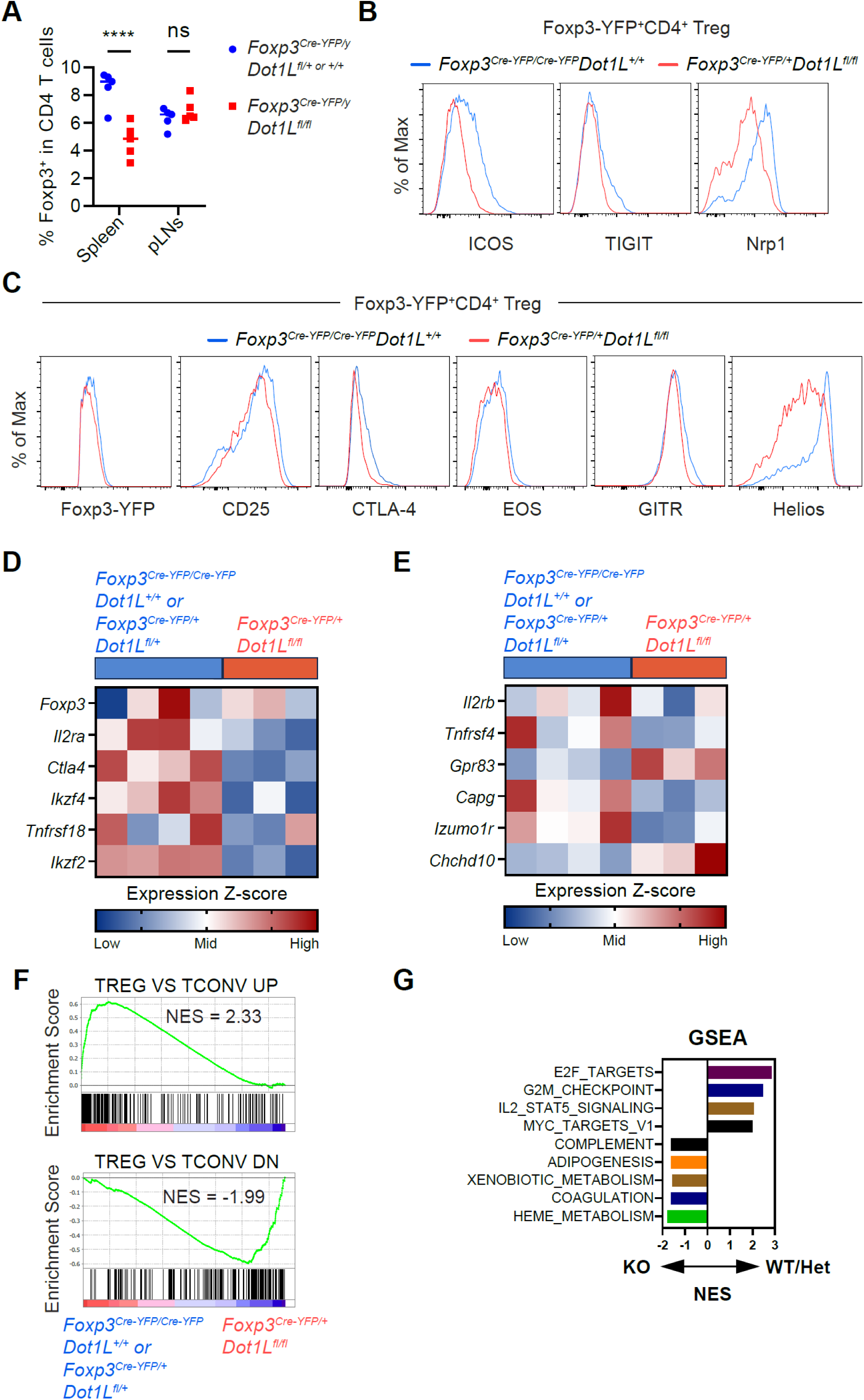
Treg-specific Dot1L deletion compromises Treg identity gene expression. (A) Frequencies of Foxp3^+^ Tregs in CD4^+^ T cells in spleens and pLNs of 3-to 5-week-old male *Foxp3^Cre-YFP/y^Dot1L^fl/fl^* and *Foxp3^Cre-YFP/y^Dot1L^fll+ or +/+^* littermate control mice. n = 5. (B,C) Flow cytometry of the expression of Treg effector molecules ICOS, TIGIT, and Nrp1 (B) and the expression of TSDR-associated gene products Foxp3-YFP, CD25, CTLA-4, EOS, GITR, and Helios (C) in Foxp3-Cre-YFP^+^ Tregs from female *Foxp3^Cre-YFP/Cre-YFP^Dot1L^+/+^* and *Foxp3^Cre-YFP/Cre-YFP^Dot1L^fl/fl^* mice. (D,E) Heatmaps showing expression of TSDR-associated genes (D) and Treg specific genes not associated with known TSDRs (E) in Foxp3-Cre-YFP^+^ Tregs from female *Foxp3^Cre-YFP/+^Dot1L^fl/fl^* and control *Foxp3^Cre-YFP/Cre-YFP^Dot1L^+/+^* or *Foxp3^Cre-YFP/+^Dot1L^fl/+^* mice. (F) Gene Set Enrichment Analysis (GSEA) of the expression of Treg-specific genes in Foxp3-Cre-YFP^+^ Tregs from female *Foxp3^Cre-YFP/+^Dot1L^fl/fl^* and control *Foxp3^Cre-YFP/Cre-YFP^Dot1L^+/+^* or *Foxp3^Cre-YFP/+^Dot1L^fl/+^* mice. (G) GSEA of the expression of indicated gene sets in Foxp3-Cre-YFP^+^ Tregs from female *Foxp3^Cre-YFP/+^Dot1L^fl/fl^* and control *Foxp3^Cre-YFP/Cre-YFP^Dot1L^+/+^* or *Foxp3^Cre-YFP/+^Dot1L^fl/+^* mice. Mean ± SEM. ****p < 0.0001, one-way ANOVA and Holm-Šídák test in (A).

Next, to define Dot1L’s impact on the global Treg transcriptome, we performed RNA-seq on CD4^+^YFP^+^ Tregs isolated from non-inflammatory *Foxp3^Cre-YFP/+^Dot1L^fl/fl^*and control littermates. Transcripts of TSDR-associated genes were significantly downregulated in Dot1L-deficient Tregs (**Fig. 6D**), consistent with the protein-level defects. In contrast, expression of core Treg genes lacking documented TSDRs remained unaltered **(Fig. 6E**), highlighting Dot1L’s selective regulation of TSDR-linked programs. Gene set enrichment analysis (GSEA) further revealed that Dot1L deletion impaired the signature of genes upregulated in Tregs versus conventional CD4^+^ T cells while enriching for genes normally repressed in Tregs (**Fig. 6F**). Pathway analysis also uncovered reductions in E2F and Myc target genes, G2/M checkpoint regulators, and IL-2/STAT5 signaling components, accompanied by upregulation of several other pathways (**Fig. 6G**). Together, these findings establish Dot1L as a critical epigenetic regulator of the Treg-specific gene expression program and suggest additional roles in pathways that support Treg homeostasis and function.

### Dot1L-dependent Treg Gene Programming Ensures Lineage Stability and Suppressive Capacity

To establish whether Dot1L-dependent DNA demethylation underlies both Treg lineage stability and in vivo suppressive function, we asked whether transient Dot1L inhibition during tTreg differentiation leads to durable impairments in Treg-specific gene expression and function, and whether enhancing TET activity with VC could rescue these defects using a standard in vivo co-transfer suppression assay (**Fig. 7A**). Pre-tTregs from CD45.2^+^*Foxp3^Thy1.1^* mice were activated with or without the Dot1L inhibitor SGC in the presence of low or high concentrations of VC. These tTregs were co-transferred together with CD45.1⁺Foxp3^DTR^ effector T cells into *Rag1*^⁻*/*⁻^ recipients. tTregs generated with low VC (no SGC) markedly reduced both weight loss and donor Teff expansion (**Fig. 7, B and C**). In contrast, transient Dot1L inhibition under low VC abolished protection—mice lost more weight and harbored higher Teff frequencies—demonstrating that Dot1L activity during differentiation is critical for tTreg function. Strikingly, augmenting VC partially rescued suppressive capacity even in the presence of SGC, indicating that boosting TET-mediated demethylation can compensate for Dot1L loss.

**Figure 7.**
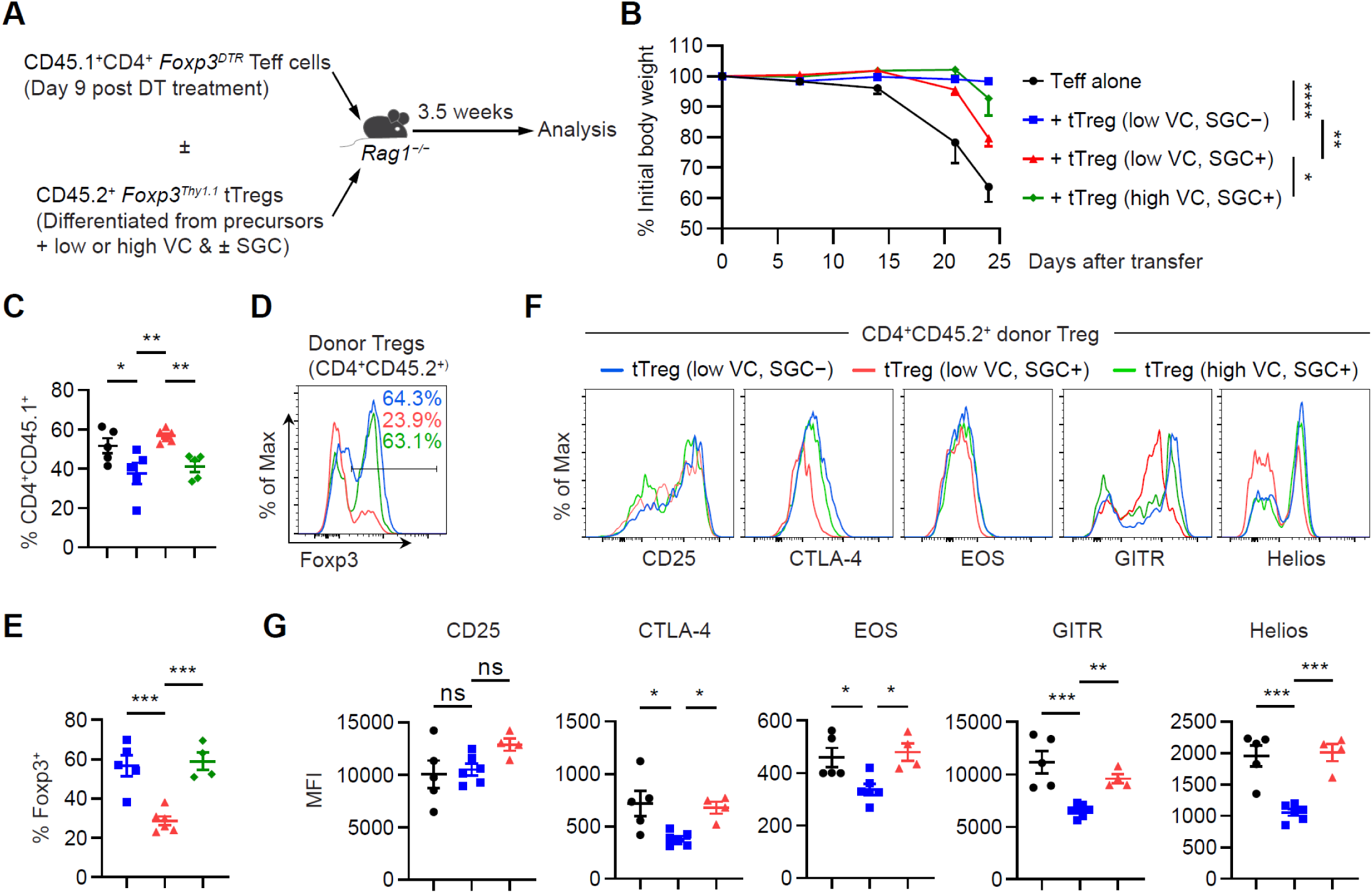
Dot1L-dependent specification of Treg gene expression is critical for Treg lineage stability and suppressive function. (A) Schematic representation of the adoptive transfer experiment for assessing tTreg lineage stability and suppressive function in vivo. CD4^+^ effector T (Teff) cells were isolated from *CD45.1^+^Foxp3^DTR^* mice at day 9 post diphtheria toxin (DT) treatment. Foxp3-Thy1.1^+^ tTregs were differentiated in vitro from *Foxp3^Thy1.1^* tTreg precursors in the presence of 0.2 µg/ml (low) or 10 µg/ml (high) of VC and with or without 2 µg/ml of SGC. n = 5-6. (B) Percentages of initial body weight of *Rag1^−/−^* recipient mice at indicated days post adoptive cell transfer. (C) Frequencies of CD4^+^CD45.1^+^ Teff cells in total living cells in *Rag1^−/−^* recipient mice 24 days after adoptive transfer. (D,E) Flow cytometry (D) of Foxp3 expression and frequencies (E) of Foxp3^+^ cells in CD45.2^+^CD4^+^ donor tTregs in *Rag1^−/−^* recipient mice 24 days after adoptive transfer. (F,G) Flow cytometry (F) and mean fluorescence intensity (MFI) (G) of CD25, CTLA-4, EOS, GITR, and Helios in donor tTregs in *Rag1^−/−^* recipient mice 24 days after adoptive transfer. Mean ± SEM. *p < 0.05, **p < 0.01, ***p < 0.001, ****p < 0.0001, one-way ANOVA and Holm-Šídák test in (B, C, E, and G).

We then assessed whether these functional differences correlated with Foxp3 stability and expression of TSDR-associated genes *in vivo*. tTregs differentiated with SGC (low VC) exhibited substantially reduced Foxp3 retention and lower levels of key Treg markers—CTLA-4, EOS, GITR, and Helios—compared to controls (**Fig. 7, D and G**). Importantly, raising VC concentration fully restored both Foxp3 stability and the expression of these TSDR-linked proteins despite Dot1L inhibition. Together, these data establish that Dot1L promotes Treg lineage stability and suppressive function by facilitating TET-driven DNA demethylation during Treg differentiation.

## Discussion

Both Treg-specific DNA demethylation and Foxp3 expression are essential for establishing the Treg gene expression program and full suppressive capacity (4). We recently demonstrated that sustained Foxp3 transcription promotes CNS2 demethylation and stabilizes Foxp3 expression in iTregs (20). However, the mechanism linking prolonged transcriptional activation to DNA demethylation—and whether it broadly governs TSDR demethylation in a gene- and cell-type-specific fashion—remains unclear.

Our findings implicate Dot1L-mediated H3K79 methylation as a key facilitator of TSDR demethylation. H3K79 methylation is enriched within gene bodies and correlates with transcriptional activity (26–28). Because most TSDRs reside in transcribed regions of genes upregulated during Treg differentiation (4), Dot1L-dependent H3K79 methylation likely provides a general mechanism by which CpG-rich enhancers within these highly induced genes are demethylated. Once demethylated and activated, these enhancers may serve as an epigenetic “memory” of prior gene activation, thereby reinforcing the expression of critical Treg-specific genes that underlie lineage stability and functional fidelity. Future studies will be needed to determine whether diminished Dot1L function contributes to defective Treg differentiation or instability in autoimmune disease and, conversely, whether enhancing Dot1L activity could be leveraged to strengthen Treg lineage stability and function for therapeutic benefit.

Our findings demonstrate that Dot1L-mediated H3K79 methylation enhances CNS2 chromatin activation, consistent with its established role at enhancers in leukemia cells (34, 35). Critically, directing dCas9-VPR to CNS2 in naïve T cells—and thereby enforcing chromatin activation—restored CNS2 demethylation even when Dot1L was inhibited, indicating that an open chromatin state is a prerequisite for downstream DNA demethylation. Additionally, Dot1L inhibition markedly reduced TET protein recruitment to CNS2, suggesting that H3K79 methylation facilitates enhancer activation and thereby promotes TET binding and DNA demethylation at this locus.

Mice deficient for CNS2 develop a late-onset, nonfatal inflammatory disorder (18, 19). In stark contrast, Treg-specific deletion of Dot1L triggers an early-onset, fatal inflammatory syndrome, accompanied by impaired DNA demethylation not only at CNS2 but across multiple TSDRs. Remarkably, enhancing TET activity with high-dose vitamin C during in vitro tTreg differentiation fully restores both TSDR-associated gene expression and suppressive function despite Dot1L inhibition. These data demonstrate that demethylation of multiple TSDRs is non-redundantly essential for Treg identity and function.

Intriguingly, Dot1L-deficient mice develop more severe disease than mice lacking Tet2 and Tet3 in Tregs (36), even though both strains exhibit comparable TSDR demethylation defects. This disparity suggests that Dot1L also supports Treg function through DNA-demethylation-independent mechanisms—most likely by promoting chromatin activation and transcription factor recruitment at gene-body enhancers. Indeed, our gene-set enrichment analysis reveals widespread disruption of Treg-specific gene expression upon Dot1L deletion, including genes lacking TSDRs but highly expressed in effector Tregs. Together, these findings position Dot1L as a multifaceted regulator of Treg cellular identity, acting through both epigenetic demethylation and direct enhancer activation.

In summary, our findings reveal a transcription-coupled mechanism in which activation of Treg-specific genes during differentiation recruits Dot1L to deposit H3K79 methylation. This modification enhances chromatin accessibility and facilitates TET enzyme binding, driving TSDR demethylation. Crucially, Dot1L-dependent demethylation and the resulting gene expression program are essential for Treg lineage stability and suppressive function—loss of which precipitates a fatal inflammatory disorder. This model of transcription-linked chromatin remodeling establishing DNA-demethylation-based epigenetic memory offers a promising strategy for therapeutic modulation of Treg identity and function.

## Materials and Methods

### Mice

Animals were housed at the Tufts University School of Medicine (TUSM) animal facility under specific pathogen-free conditions according to institutional guidelines. All studies were performed under protocol B2022-85 and approved by Tufts Institutional Animal Care and Use Committee. All mouse strains used were on the C57BL/6 genetic background. B6 CD45.1 (#002014), B6 CD45.2 (#000664), *Foxp3^Cre-YFP^* (#016959), *Foxp3^DTR^* (#016958), *Rag1^−/−^* (#002216), and *Rosa^Cas9-eGFP^* (#026179) mice were purchased from the Jackson Laboratory. *Foxp3^Thy1.1^* mice were a gift from Dr. Y. Zheng. *Dot1L^fl/fl^* mice were a gift from Dr. Piroska Szabo. The above mouse strains were bred in-house at TUSM to produce the *Foxp3^Thy1.1^ Rosa26^Cas9^*, *Foxp3^Cre-YFP^Dot1L^fl/fl^*, *Foxp3^Cre-YFP^Dot1L^+/+^* or *Foxp3^Cre-YFP/+^Dot1L^fl/+^*, CD45.1^+/+^ *Foxp3^DTR^* strains.

### Reagents and flow cytometry

Anti-CD3 and anti-CD28 were from BioXCell. Anti-CD3/CD28 beads were from Miltenyi Biotec. Human IL-2 was from PeproTech. Mouse TGF-b and IL-6 were from R&D Systems. Vitamin C, Diphtheria toxin (DT), and SGC0946 were obtained from Sigma-Aldrich. Abs for flow cytometry were from BioLegend, eBioscience, and Tonbo Biosciences. After preparation of single-cell suspensions red-blood cell lysis was preformed and cell counts were determined using hemacytometer. For surface staining, cells were stained for 15 min with fluorochrome-conjugated antibodies before washing and analysis or intracellular staining. For intracellular staining, cells were fixed and permeabilized using the Foxp3 Transcription Factor Staining Buffer Kit (Tonbo Biosciences) followed by staining with fluorochrome-conjugated antibodies diluted in Perm/Wash buffer. Flow cytometry was performed using the LSRII (BD Biosciences) and data were analyzed with FlowJo software (Tree Star).

For cytokine staining single-cell suspensions were stimulated with PMA (50 ng/ml) and Ionomycin (500 ng/ml) for 2 h at 37°C followed by 2 h incubation with Golgi-Stop (1:1500 dilution, BD Biosciences) at 37°C. Cells were spun down and stained using intracellular staining protocol described above.

### Retroviral vectors and transduction

Single-guide RNA (sgRNA) vector MG2N was generated by modifying MSCV-P2GM-FF (Addgene). sgRNA were cloned into BbsI-digested MG2N. The guide sequences are non-targeting (sgNT) (5’-GCACTACCAGAGCTAACTCA-3’), Dot1L knockout (sgDot1L) (5’-CTGCAAACATCACTACGGAG-3’). CNS2 targeting (sgCNS2) (5’-GGGCTTCATCGGCAACAAGG-3’) Vector MdC-VPRC (dCas9-VPR), was generated by replacing IRES-eGFP on MIGR1 (Addgene) with expression cassette (dCas9-VPR-P2A-hCD2).

For Cas9 mediated knockout of Dot1L, Tn cells from *Rosa^Cas9-EGFP^Foxp3^Thy1.1^* mice were purified with Naive CD4^+^ T Cell Isolation Kit (Miltenyi Biotec) and stimulated for 1 day with plate-coated anti-CD3 (5 µg/ml) and anti-CD28 (5 µg/ml) in the presence 200 U/ml IL-2, followed by spin infection with MG2N-sgRNA vector at in the presence of 4 µg/ml polybrene. Post-transduction media was supplemented with 1000 U/ml IL-2 and 2 ng/ml TGF-β. Five days post-plating CD4+ NGFR^+^ Thy1.1^+^ cells were sort-purified and re-stimulated with anti-CD3/CD28 beads (2:1 beads/cells ratio) in the presence of 1000 U/ml IL-2, 0.5 ng/ml TGF-β and 1 µg/ml vitamin C.

For targeting dCas9-VPR to CNS2 in UT or SGC treated Tconv cells, Tn cells from Foxp3^Thy1.1^ mice were purified with Naive CD4^+^ T Cell Isolation Kit (Miltenyi Biotec) and stimulated for 1 day with plate-coated anti-CD3 (5 µg/ml) and anti-CD28 (5 µg/ml) in the presence 200 U/ml IL-2, followed by spin infection at in the presence of 4 µg/ml polybrene. Post-transduction media was supplemented with 1000 U/ml IL-2. Five days post-plating CD4^+^ NGFR^+^ hCD2^+^ cells were sort-purified and re-stimulated with anti-CD3/CD28 beads (2:1 beads/cells ratio) in the presence of 1000 U/ml IL-2 and 1 µg/ml vitamin C.

### Generation of SGC-treated iTregs/RSiTregs

Tn cells from *Foxp3^Thy1.1^* mice were purified with Naive CD4^+^ T Cell Isolation Kit (Miltenyi Biotec) and stimulated for 1 day with plate-coated anti-CD3 (5 µg/ml) and anti-CD28 (5 µg/ml) in the presence 200 U/ml IL-2, 2 ng/ml TGF-β, and presence or absence of 2 μM SGC0946. After 4 days cells were re-stimulated with anti-CD3/CD28 beads (2:1 beads/cells ratio) in the presence of 1000 U/ml IL-2, 0.5 ng/ml TGF-β, 1 µg/ml vitamin C, and presence or absence of 2 μM SGC0946. At experimental endpoint Foxp3-Thy1.1^+^ cells were column purified using anti-PE MicroBeads (Miltenyi Biotec) for bisulfite sequencing.

In assays examining ability of VC to rescue Dot1L dependent inhibition of CNS2 demethylation, 2 µg/ml or 10 µg/ml of VC was used during iTreg differentiation with or without 2 μM SGC0946.

### In vitro Foxp3 stability assays

For in vitro Foxp3 stability assay, Foxp3-Thy1.1^+^ cells were column purified using anti-PE MicroBeads (Miltenyi Biotec), washed and then replated with 100 U/ml IL-2 and 20 ng/ml of IL-6 (R&D systems).

### In vivo CNS2 demethylation assay

Foxp3Thy1.1^+^ iTregs doubly transduced with Cas9-RV and MG2N targeting Dot1L or the control non-targeting vector were co-transferred with congenically distinct Tn cells into *Rag1^−/−^* mice via retro-orbital injection. Five weeks post-injection, Foxp3Thy1.1^+^ donor iTregs were sort-purified for bisulfite sequencing.

### Bisulfite sequencing

Genomic DNA was bisulfite converted using EpiTect Bisulfite Conversion Kit (Qiagen), amplified with Q5U polymerase (New England Biolabs) using previously described primers listed below. Amplified DNA is purified after agarose gel electrophoresis and cloned into pJET1.2 (Thermo Fisher Scientific) for Sanger sequencing.

CNS2-F (5’-TGGGTTTTTTTGGTATTTAAGAAAG-3’)

CNS2-R (5’-AACCAACCAACTTCCTACACTATCTAT-3’)

Ctla4-F (5’-TGGTGTTGGTTAGTAGTTATGGTGT-3’)

Ctla4-R (5’-AAATTCCACCTTACAAAAATACAATC-3’)

IL2a-F (5’-TTTTAGAGTTAGAAGATAGAAGGTATGGAA-3’)

IL2ra-R (5’-TCCCAATACTTAACAAAACCACATAT-3’)

Ikzf2-F (5’-AGGATGGTTTTTATTGAAGGTGAT-3’)

Ikzf2-R (5’-GGTGTAGTGTTTGTTTGGTGTGTAT-3’)

Ikzf4-F (5’-TAAGAAATTGGGTGTGGTATATGTA-3’)

Ikzf4-R (5’-AAGGAGTTTTAGTAGGGGAAA-3’)

Tnfrsf18-F (5’-GAGGTGTAGTTGTTAGTTGAGGATGT-3’)

Tnfrsf18-R (5’-AACCCCTACTCTCACCAAAAATATAA-3’)

### Chromatin immunoprecipitation and qPCR

Samples for chromatin immunoprecipitation were prepared using SimpleChIP Plus enzymatic chromatin IP kit (Cell Signaling 9005) according to manufacturer instructions. Briefly, 1e6 cells were crosslinked in media containing 1% formaldehyde for 10 min. Crosslinking was quenched with addition of glycine (final concentration 1x). Crosslinked cells were lysed, and nuclei were pelleted via centrifugation before digestion with micrococcal nuclease. After digestion samples were sonicated to shear nucleosomal DNA. Samples were centrifuged and chromatin-containing supernatant was removed. Prior to immunoprecipitation, aliquot of chromatin-containing supernatant was set aside as input control. Samples were then aliquoted for individual chromatin immunoprecipitation reactions, with 1 μl of desired antibody used in each reaction. Samples were incubated overnight at 4°C with rotation. Bead-based purification was used to precipitate antibody-bound complexes followed by reversal of cross-linking and DNA precipitation. Relative abundance of immunoprecipitated DNA fragments were assessed by qPCR on a QuantStudio 6 Flex instrument with region specific primers listed below. Enrichment was calculated relative to total input.

CNS0-F (5’-AGCAGCACACAGGCCTTAAAAC-3’)

CNS0-R (5’-GGTTTTCTTTGTAGGTCGGTGAC-3’)

*Foxp3* TSS-F (5’-GCAGTTTCCCACAAGCCAGGCT-3’)

*Foxp3* TSS-R (5’-GGAGAGCAGGGACACTCGCTCA-3’)

CNS1-F (5’-CTGTGCATGGGTCTCTGCCACG-3’)

CNS1-R (5’-GCCAGAGACACCCATGGCTGGT-3’)

CNS2-F (5’-GGGCCCAGATGTAGACCCCGAT-3’)

CNS2-R (5’-CCAGCCAGCTTCCTGCACTGTC-3’)

CNS3-F (5’-TCTCCAGGCTTCAGAGATTCAAGG-3’)

CNS3-R (5’-ACAGTGGGATGAGGATACATGGCT-3’)

ChIP antibodies purchased from Cell Signaling include H3K27ac (#8173), H3K9ac (#9649), H3K27me3 (#9733), H3K4me3 (#9751), H3K4me1 (#5326), H3K79me2 (#5427), and Tet2 (#92529). Tet3 (#ABE290) antibody was purchased from MilliporeSigma.

### Isolation and culture of thymic Tregs and Treg precursors

Thymi were isolated from *Foxp3^YFP-Cre^Dot1L^fl/fl^* and control *Foxp3^YFP-Cre^ Dot1L^+/fl^* mice at 2-3 weeks of age. After preparation of single-cell suspensions, cells were treated with anti-FcRγIII/II antibody before depletion of PE-labeled CD8^+^ T cells. The negative fraction was stained with fluorochrome-conjugated anti-mouse antibodies and subsets of CD4SP thymocytes were sort purified. For bisulfite sequencing of thymic Tregs, Foxp3-YFP^+^CD25^+^CD24^lo^CD73^hi^ cells were sort-purified for genomic DNA extraction and bisulfite sequencing. CD4^+^Thy1.1^−^CD25^hi^CD73^lo^thymic Treg precursors (pre-tTreg) cells were sort-purified from 11-day old *Foxp3^Thy1.1^* mice and stimulated with anti-CD3/CD28 beads in the presence of 1000 U/ml IL-2 with or without 2 μM SGC for six days before extracting genomic DNA for bisulfite sequencing.

### RNA-sequencing and analysis

Single-cell suspensions were generated from pooled spleen and lymph nodes isolated from *Foxp3^YFP-Cre^ Dot1L^fl/fl^* and control *Foxp3^YFP-Cre^ Dot1L^+/+^* female mice. Total CD4^+^ cells were isolated with CD4^+^ T Cell Isolation Kit (Miltenyi Biotec). The positive fraction was stained with fluorochrome-conjugated anti-mouse antibodies for sort-purification of CD4^+^ Foxp3-YFP^+^ Tregs using FACSAria II to 99% purity. Total RNA was extracted using TRIzol and unstranded cDNA libraries were prepared. RNA was isolated using phenol-chloroform extraction. Uniquely indexed libraries were pooled in equimolar ratios and sequenced on a single Illumina NextSeq500 run with single-end 75-bp reads by the Tufts University Genomics Core Facilities.

Sequence reads were aligned with the mm39 reference genome assembly and gene counts were quantified with FeatureCounts. Differential expression analysis was performed with DESeq2. Gene-set enrichment analyses were performed with GSEAPreranked, in which genes ranked according to their fold changes were compared with the hallmark gene sets.

### Isolation of CD4^+^ Teff cells for in vivo suppression assays

*Foxp3^DTR^* mice were treated with 1 μg of diphtheria toxin (DT) via intraperitoneal injection on day 0 and day 1 and every other day thereafter. After 9 days animals were sacrificed, and single cell suspensions were prepared from pooled lymph nodes and spleen. CD4^+^ cells were isolated with CD4^+^ T Cell Isolation Kit (Miltenyi Biotec). The positive fraction was stained with fluorochrome-conjugated anti-mouse antibodies for sort-purification of CD4^+^ CD44^hi^ CD62L^lo^ effector T cells.

### In vivo thymic Treg suppression assay

Thymic Treg precursors (CD4^+^Thy1.1^−^CD25^hi^CD73^lo^) were isolated from *Foxp3^Thy1.1^* mice and cultured in presence of 0.2 µg/ml (low) or 10 µg/ml (high) VC with or without 2 μM SGC. Thy1.1^+^ Tregs were then sort-purified and co-transferred with sort-purified CD4^+^ CD44^hi^ CD62L^lo^ effector T cells isolated from *Foxp3^DTR^* mice previously immunized with DT into *Rag1^−/−^*recipients. Post-transfer bodyweights were monitored, and animals were sacrificed on day 24 and single-cell suspensions were prepared from pooled lymph nodes for flow cytometry analysis.

### Statistics

Except for RNA-Seq analysis, statistical significance was determined using GraphPad Prism 10.0 (GraphPad Software). For comparisons of a single variable between 2 groups, significance was determined by 2-tailed Student’s t tests. For comparisons of multiple groups where variance did not significantly differ across groups, 1- or 2-way ANOVA with Šidák’s (for comparisons between preselected pairs) multiple-comparison corrections was used. P values below 0.05 were considered statistically significant and are shown by the exact number or by asterisks in the figures.

## Study approval

All studies were performed under protocol B2022-85 and approved by Tufts Institutional Animal Care and Use Committee.

## Data and materials availability

All data are available in the main text. RNA-Seq data can be accessed using GEO accession no. GSE313284. All materials used or generated in this study are available to researchers following appropriate standard material transfer agreements.

## Author contributions

Conceptualization: X.L. and J.J.C.

Methodology: X.L. and J.J.C.

Investigation: J.J.C. T.R.C., and X.L.

Funding acquisition: X.L.

Project administration: X.L.

Supervision: X.L.

Writing: J.J.C. and X.L.

## Acknowledgments

X.L. was supported by the Ellison Foundation, the NIH grant AI192616 (awarded to X.L.), and the NIH grant AI167245 (awarded to A.N.P.). J.J.C. was supported by a Tufts University Graduate School of Biomedical Sciences Dean’s Fellowship. T.R.C. was supported by the NIH grant AI167245 (awarded to A.N.P.). We thank Y. Zheng and A. Rudensky for sharing various mouse strains for this study; S. Kwok and A. Parmelee for assistance with FACS sorting; A. Tai and I. Grinvald for assistance with RNA-seq experiments; P. Alcaide, C. Genco, and M. Gaglia for helpful discussions.

